# Maintaining maximal metabolic flux by gene expression control

**DOI:** 10.1101/115428

**Authors:** Robert Planqué, Josephus Hulshof, Bas Teusink, Johan Hendriks, Frank J. Bruggeman

## Abstract

Many evolutionarily successful bacteria attain high growth rates across growth-permissive conditions. They express metabolic networks that synthesise all cellular components at a high rate. Metabolic reaction rates are bounded by the concentration of the catalysing enzymes and cells have finite resources available for enzyme synthesis. Therefore, bacteria that grow fast should express needed metabolic enzymes at precisely tuned concentrations. To maintain fast growth in a dynamic environment, cells should adjust gene expression of metabolic enzymes. The activity of many of the associated transcription factors is regulated by their binding to intracellular metabolites. We study optimal metabolite-mediated regulation of metabolic-gene expression that preserves maximisation of metabolic fluxes across varying conditions. We logically derive the underlying control logic of this type of optimal regulation, which we term ‘Specific Flux (*q*) Optimization by Robust Adaptive Control’ (*q*ORAC), and illustrate it with several examples. We show that optimal metabolic flux can be maintained in the face of ***K*** changing parameters only if the number of transcription-factor-binding metabolites is at least equal to ***K***. *q*ORAC-regulation of metabolism can generally be achieved with basic biochemical interactions, indicating that metabolism can operate close to optimality. The theory that we present is directly applicable to synthetic biology, biotechnology and fundamental studies of the regulation of metabolism.

## Highlights

- A general framework called *q*ORAC is presented which dynamically steers a given metabolic pathway to maximal steady state specific flux in changing external conditions, and which follows logically from the assumptions.
- We show that maximising specific flux has a unique solution for a large class of enzymatic rate laws.
- The *q*ORAC framework uses metabolite-binding transcription factors to influence gene expression. The metabolites are called sensors.
- The metabolic pathway may be made robust to changes in *K* parameters if there are *K* sensors to control gene expression.

## Introduction

Microbes need to grow fast to outcompete others, and therefore need to maintain high growth rates in changing environments. Their specific fluxes (metabolic rates per unit of expended enzyme) thus need to be kept maximal. Since catalytic enzymes are a limited resource, this requires cells to be economical, synthesise the right enzymes, in the right amounts, and adapt to fluctuating nutrient levels.

Experimental evidence is mounting that cells are indeed able to tune enzyme levels to maximise the growth rate (Figure 1; Walsh and Koshland (1985);van der Vlag et al. (1994);Jensen et al. (1995);Andersen et al. (2001);Solem et al. (2003);Koebmann et al. (2005);Dekel and Alon (2005);Keren et al. (2016)). Efficient enzyme allocation has also recently been shown to underlie a surprising number of other general physiological phenomena such as the bacterial growth laws (Scott et al., 2010, 2014;Bosdriesz et al., 2015), overflow metabolism (the Crabtree or Warburg effect;Molenaar et al. (2009);Basan et al. (2015);Weiße et al. (2015)), and catabolite repression (You et al., 2013). Except perhaps for the case of optimal ribosomal synthesis (Scott et al., 2014;Bosdriesz et al., 2015), it is not clear in any of these examples how cells achieve the optimisation, i.e., how to find a new optimum when conditions have changed.

**Figure 1.**
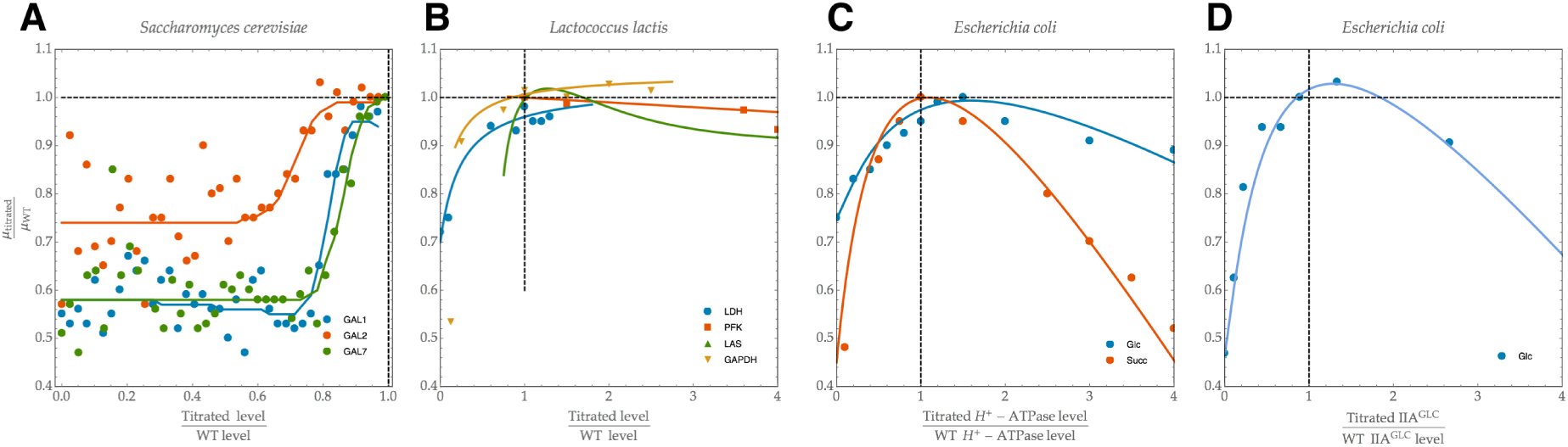
Experimental evidence indicating that microbes tune their enzyme levels to maximise growth rate. In each example, the wild type (WT) is shown to express enzyme concentrations at which the growth rate *μ* is approximately maximal. Data adapted from: A,Keren et al. (2016); B,Andersen et al. (2001);Solem et al. (2003);Koebmann et al. (2005), C,Jensen et al. (1995); D, (van der Vlag et al., 1994). Abbreviations: GAL1, galactokinase; GAL2, Galactose permease; GAL7, Galactose-1-phosphate uridyl transferase; LDH, lactate dehydrogenase; PFK, phosphofructokinase; LAS, las operon; GAPDH, glyceraldehyde 3-phosphate dehydrogenase; Glc, glucose; Succ, succinate. In (Keren et al., 2016) there are many other examples, including several proteins that do not show levels at which growth rate is optimised.

Gene expression regulation is largely achieved by transcription factors that are either affected by signal transduction cascades, or by direct binding of metabolites, as readouts of cellular states. Even though the latter mode of regulation, using metabolites as sensors (Kotte et al., 2010;Kochanowski et al., 2013), is widely accepted in the field, the identity of such internal cues is only known in a handful of cases (Figure 2). In *E. coli,* fructose-1,6-bisphosphate (FBP), a glycolytic intermediate, binds to the transcription factor Cra to regulate genes involved in glycolysis (You et al., 2013;Kotte et al., 2014); in yeast, the galactose catabolic pathway is induced by intracellular galactose (Sellick et al., 2008); uncharged tRNAs bind to ppGpp to mediate amino acid metabolism and ribosome synthesis (Scott et al., 2014;Bosdriesz et al., 2015); the amino acid L-Tryptophan regulates the transcription of several enzymes involved in its biosynthetic pathway (Gollnick et al., 2005); perhaps the best known example is the lactose operon, which is induced by allolactose, an intermediate of the pathway (Gilbert and Müller-Hill, 1966). There is even very recent experimental evidence that *E. coli’s* central metabolism is in fact controlled by just three such sensor metabolites (cyclic AMP (cAMP), FBP and fructose-1-phosphate (F1P);Kochanowski et al. (2017)).

**Figure 2.**
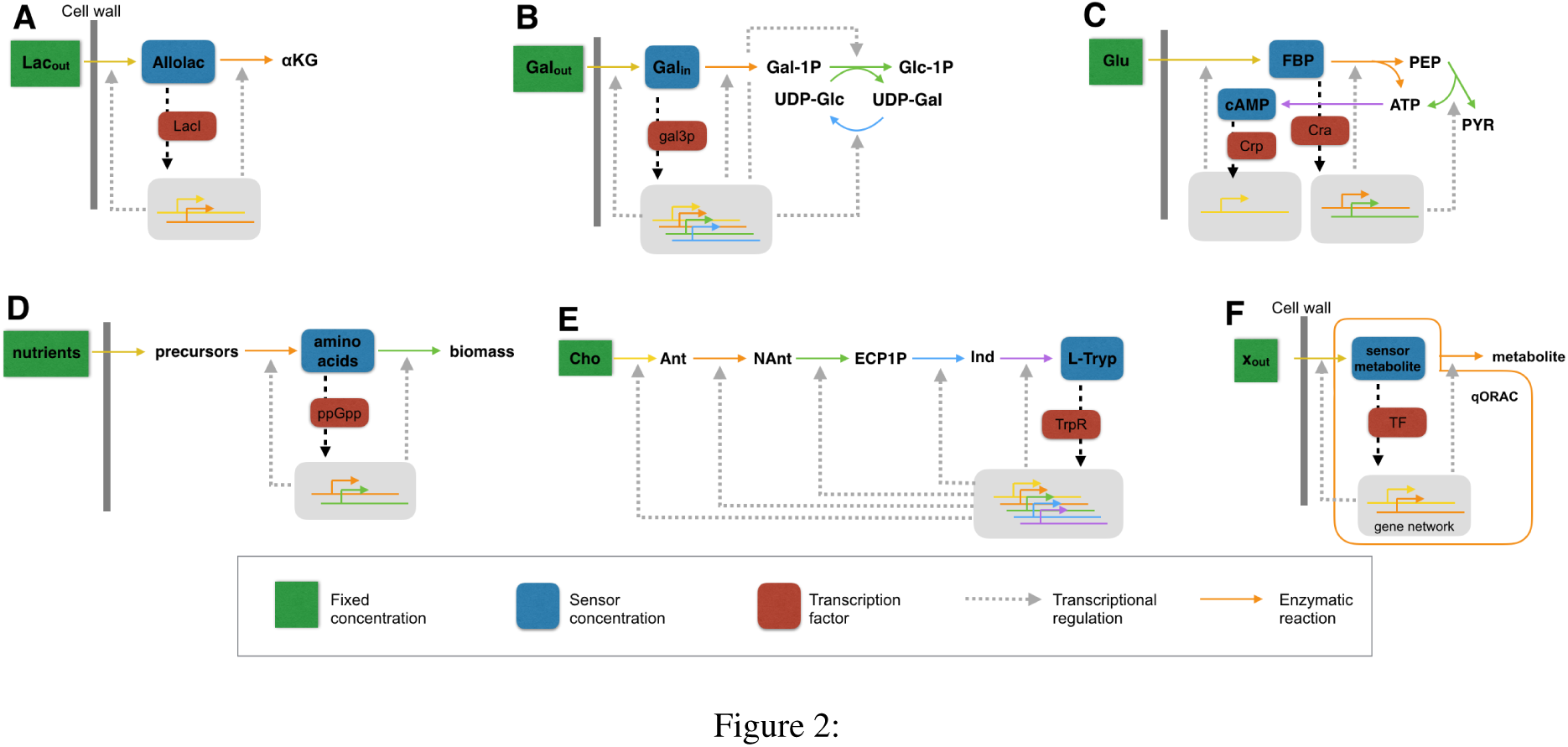
Biological examples of *q*ORAC. Five well-characterised metabolic pathways in which a metabolite binds to a transcription factor (TF) to influence gene expression. The *q*ORAC framework applies to each of them: in each case, the *q*ORAC formalism gives rise to the enzyme synthesis rates that steer the metabolic pathway to maximal metabolic rates that are robust to changes in the external concentration (external with respect to the pathway). (A) The lac operon in *E. coli*, with sensor Allolactose binding to LacI; (B) The galactose uptake system in yeast, with sensor internal galactose binding to gal3p; (C) The control of glycolytic enzymes via sensors FBP (binding to Cra), and cAMP (binding to Crp); (D) The control of catabolism vs biomass synthesis, with unsaturated tRNAs (due to a lack of amino acids) binding to ppGpp; (E) The control of the L-Tryptophan biosynthesis pathway by the amino acid binding to TrpR; (F) The general scheme of a *q*ORAC-steered pathway. Abbreviations: Lac_ou_t, external lactose; Allolac, allolactose; αKG, *α-* ketoglutarate; Gal_out_, external galactose; Gal_in_, internal galactose; Gal-1P, galactose-1-phosphate; Glc-1P, glucose-1-phosphate; UDP-Glc, uridine-diphosphate-glucose; UDP-Gal, uridine-diphosphate-galactose; Glu, glucose; FBP, fructose-1,6-biphosphate; PEP, phosphoenolpyruvate; PYR, pyruvate; cAMP, cyclic AMP; ATP, adenosine-triphosphate; ppGpp, guanosine tetraphosphate; Cho, chorismate; Ant, Anthranilate; NAnt, N-(5’-phosphoribosyl)-anthranilate; ECP1P, Enol-1-0-carboxy-phenylamino-1-deoxyribulose phosphate; Ind, Indole-3-glycerol-P; L-Tryp, L-tryptophan.

What has remained unexplained so far is why certain sensor metabolites play this role and others do not, why there are just a few of them, and more importantly, what type of enzyme synthesis rates the gene network must produce in order to steer a metabolic pathway in the direction of maximal specific flux when conditions change.

We will show that all three problems admit a general answer by deriving one unified framework, which we call *q*ORAC (for Specific Flux (*q*) Optimisation by Robust Adaptive Control). We will show that for practically any given pathway, it is possible to derive the enzyme synthesis rates which the gene network must produce to steer the pathway to its optimum, on the basis of internal metabolic sensor information alone. We moreover show that it is possible to make predictions on the identity of metabolite sensors, and we prove that a pathway that is robust to changes in *K* external or internal parameters must be influenced by (at least) *K* sensor metabolites. The control dynamics follow logically from the assumptions and are thus in that sense unique. They do not depend on the species, kinetics, allosteric control or parameter settings; the actual implementation of the control dynamics of course does.

We will discuss how all of the current known examples of metabolites acting as sensors cited above may be interpreted within the *q*ORAC framework. We thereby show that it is indeed possible for gene networks to steer given pathways dynamically and robustly to high specific fluxes. The theory thus shows that surprising recent findings such as the pervasive optimisation of enzyme levels in yeast (Keren et al., 2016), or the small number of sensor metabolites found in *E. coli’s* central metabolism (Kochanowski et al., 2017), are in fact to be expected.

## Results

### A biological example: the galactose pathway in yeast

We will first explain the cellular control problem that we will solve, by considering the well-understood biological example of the galactose uptake system in yeast (Figures 1A, 2B). We wish to maximise the steady state flux through this pathway. A finite pool of enzymes has to be distributed across the four different enzymatic reactions to optimise metabolic flux. Depending on the external galactose concentration, more or less enzyme should be invested in the first transporter step, from Gal_out_ to Gal_in_. This leaves a correspondingly smaller or larger pool of enzymes for the rest of the pathway. Keeping enzyme concentrations fixed for the moment, increasing Gal_out_ will cause Gal_in_ to increase as well, so this should be indicative for a change in conditions and thus a signal for the adaptation of enzyme concentrations: the transporter enzyme concentration should decrease, and the other enzymes should increase con-comittantly.

The question is now how to find the optimal allocation of enzyme concentrations after such a change, just on the basis of the value Gal_in_. Gal_in_ plays the role of metabolic sensor, relaying information to the gene network by binding to a transcription factor, in this case gal3p. The gene network should then induce the right genes at the right rate to change the enzyme concentrations in the pathway such that they finally reach their optimal steady state level, at which the specific flux through the pathway is maximal.

Having found optimal input-output relations for the gene circuit, with concentrations of sensor metabolites as input and enzyme concentrations as output, the question remains whether a gene network may be found that can implement those relations. Since the gene network for the galactose pathway in yeast is known, this may be done by fitting parameters in this network (Berkhout et al., 2013). In this paper, however, we show that the problem of finding optimal input-output relations for a given metabolic pathway has a general solution, and may be generated for quite arbitrary pathways such as the ones shown in Figure 2A-E.

### Optimal specific flux in a pathway: steady-state input-output relationships

We consider the dynamics of internal metabolic concentrations *x* = (*x*_i_,…*,x_n_*) in a metabolic network,

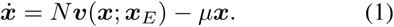

Here, *N* is the stoichiometry matrix, *υ* (*x; x_E_*) are the reaction fluxes, *x_E_* are fixed external concentrations, and *μ* is the growth rate of the culture. It is generally assumed that the dilution rate of concentrations by growth, −*μx,* is negligible for metabolism. We take the same view here, and consider

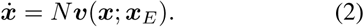

We aim to maximise the specific flux through this network, by maximising *υ_r_/e_T_*, in steady state, where *υ_r_* is some chosen output flux. Mathematically, the optimisation problem we study is

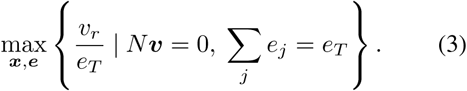

In words, we wish to maximise a given output flux per unit of total expended enzyme necessary to sustain this flux through the entire network, at steady state. This is equivalent to minimising the amount of enzyme necessary to sustain a given steady state flux *υ_r_ = V_r_*,

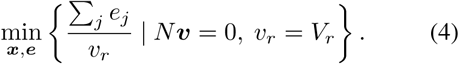

Since reaction functions generally are of the form *υ_j_ = e_j_ f_j_* (*x; x_E_*) (Cornish-Bowden, 1995), we may prescribe *υ_r_* = 1 (after all, if we can solve that problem then we can solve it for *υ_r_ = V_r_* as well by multiplying all the enzyme concentrations by *V_r_*). This allows us to rewrite (**4**) to

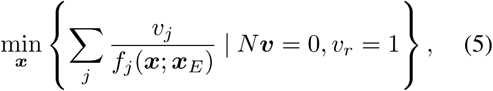

Observe that the enzyme concentration vector *e* has disappeared from the problem. It has recently been shown that the flux profiles that solve (**5**) (and therefore also the original problem (**3**)) are always subnetworks with a particularly simple structure, called Elementary Flux Modes (EFMs;Wortel et al., 2014; Müller et al., 2014). Such EFMs are one-degree-of-freedom flux vectors satisfying *Nυ = 0* that cannot be simplified further by deleting reactions without violating the steady state assumption (Schuster and Hilgetag, 1994;Schuster et al., 2002). A given EFM is thus characterised by *λ*(*V*_1_*,…,V_m_*), where *λ* is a free parameter and the flux vector (*V*_1_*,…,V_m_)* is fixed. If we want to optimise specific flux *within a given EFM* with flux vector *(V*_1_*…, V_m_*), we still need to find a vector *x* for

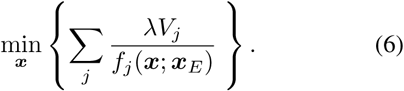

This motivates the introduction of the *objective function*

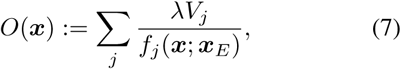

which is to be minimised, for given external concentrations *x_E_*, by suitably choosing internal concentrations *x*. This function is convex for pathways with many kinds of reaction kinetics (Liebermeister & Noor, 2015, unpublished, http://de.arxiv.org/abs/1501.02454v1), and in the SI we show that the optimum is in fact unique for an even larger class of rate laws (see also Figure S4). The objective function has a lower value if the values of *f_j_* (*x; x_E_*) are higher. Maximising specific flux may thus be reinterpreted as maximising the value of *f_j_*, which is essentially the saturation level of enzyme *j*, for all enzymes simultaneously. This can be done by making as little enzyme as possible, so that the enzymes are used at their maximal capacity.

If we find the vector *x^o^* which minimises *O*(*x*), then we can infer the corresponding optimal enzyme concentrations *e^o^* by setting

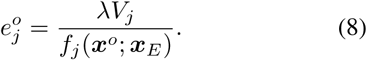

It is clear that we may choose *λ* = 1 in *O*(*x*): having found the minimiser of *O*(*x*) for *λ* = 1, we have found it for all *λ*: the corresponding enzyme levels 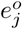 just scale with λ. In hindsight, we may also for instance normalise the enzyme concentrations such that they sum to total concentration *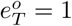*.

### Using sensors to find the optimal enzyme concentrations

The discussion so far makes clear that the optimal enzyme concentrations that maximise specific flux are defined in terms of external concentrations *x_E_*: for each choice of *x_E_*, the objective function (**7**) needs to be minimised to find *x^o^*, and subsequently *e^o^*. If we wish to design or construct gene regulatory networks that produce the right concentrations of enzymes in steady state, robustly with respect to changes in external concentrations but without direct knowledge of those changes, we see this cannot yet be done. As discussed in the Introduction, we now focus on the situation where the gene network relies only on internal information to steer metabolic gene expression. Let us thus consider a set of internal sensor metabolites, which should play the role of relaying dynamic information about changes in the environment to the gene network, such as Gal_in_ does to gal3p.

The design criterion for this gene network is that it should induce optimal enzyme synthesis rates in steady state for a whole range of external concentrations, just on the basis of sensor values. The crucial point is now that the gene network must therefore *assume* that current sensor values are in steady state and have optimal values, even when in fact they are not and are still dynamically changing. On the basis of these sensor values, the gene network then should predict the optimal steady-state enzyme levels, and causes enzyme levels to change accordingly. The immediate conclusion of this construction is that if the pathway is indeed in steady state, then so are the sensor values, and the gene network necessarily gives rise to optimal enzyme concentrations. Hence, in all likelihood, the pathway is then also in an optimal state.

Let us now construct this estimated optimum on the basis of sensor values. Steady-state optimisers *x^o^* may be characterised as minima of *O*(*x*), and are dependent on *x_E_*. Hence, *x^o^* is a critical point of *O*(*x*) *= O*(*x_i_,…,x_n_*), satisfying

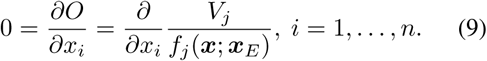

Solving (**9**) means prescribing *x_E_* and solving for *x^o^.* However, if we can solve (**9**) by prescribing some sensor values, *x_S_*, but now solving not only for the remaining internal concentrations but also for the (now unknown) external concentrations, we have found the *optimum as predicted by the sensor values, under the assumption that the pathway is in steady state.* We will denote this *predicted optimum* by *ξ* (*x_s_*). With *ξ* (*x_s_*), we can define corresponding predicted optimal enzyme levels, analogous to ( **8**),

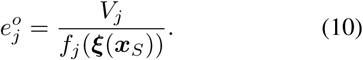

Hence, if we construct a gene network that gives rise to these optimal enzyme concentrations at steady state, we have made a system which automatically reaches the optimum if it is in steady state. The above construction is logically the *only* construction that allows for robust adaptive control, given the design criterion we have demanded the gene network to fulfil.

A simple dynamic implementation of this method can be constructed when we assume that the enzyme dynamics is described by the difference in the rates of enzyme synthesis and dilution,

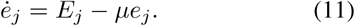

Since the enzyme synthesis rates are generally much lower than metabolic rates, we can not neglect dilution by growth in their differential equation. We assume that the cell is growing at a fixed rate μ which is minimally affected by changes in the enzyme levels that optimise specific flux of our desired EFM, and that enzyme degradation is negligible. Then, by setting

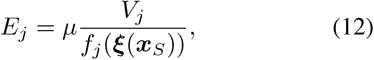

we have ensured that at steady state the enzyme levels are optimal. Note that the construction does not ensure that the combined dynamical system actually converges to this optimal steady state. The complete construction is termed *q*ORAC, and is summarised in Box 1. A simple example of a small linear pathway is specified in full detail in the SI.

##### Box 1 (*q***ORAC**)

The following differential-algebraic system of equations implements Specific Flux (*q*) Optimisation by Robust Adaptive Control (*q*ORAC) through an EFM with flux vector (*V*_1_*,…,V_m_*) in a cell culture growing at growth rate *μ*. Let *I* be the index set of internal metabolite concentrations, *E* the index set of external concentrations, and *S* the index set of sensor concentrations. Then we consider

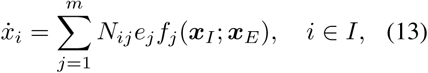

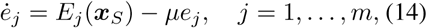

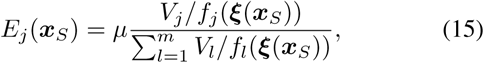

where *ξ*(*x_S_*) is the predicted optimum, and is the (time-dependent) solution of

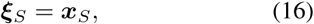

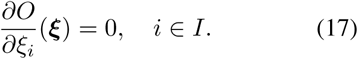

The rescaling of *E_j_* (*x_S_*) in (**15**) by the sum of all the inverses of 1/*f_j_* implies that total enzyme concentration is chosen to be equal to 1. Other rescalings give identical results, up to the chosen scaling factor. The choice above, however, is particularly useful, since it produces positive synthesis rates both for positive and negative metabolic rates through the pathway, and it ensures that it is well-defined also at thermodynamic equilibrium (see SI for details).

### The number of sensors equals the number of external parameters

The *q*ORAC construction assumes that it is possible to find internal sensor metabolites that allow a reconstruction of the optimum for different values of external concentrations *x_E_*: we need to solve (**9**) by prescribing *x_S_* and finding all the remaining variables (including the external ones). The solution for *∂O* (*x*)/ *∂x_i_ =* 0 for fixed sensor values is well-defined mathematically if the Implicit Function Theorem (IFT) holds (see SI for a more detailed exposition). Any choice of sensor metabolites for which the IFT holds is a candidate for the proposed adaptive control. An immediate consequence of the IFT is that the number of sensor metabolite concentrations must equal the number of changing external metabolite concentrations to which the system needs to be robust. This makes intuitive sense: to track changes (and hence achieve robustness) in *K* external metabolites, the gene network should be influenced by (at least) *K* (independent) internal sensors. Note also that we may not only substitute external concentrations by sensors: we could also have substituted internal parameters, such as temperature, pH, or *K_m_* values in rate laws, to make the system robust with respect to them.

### Examples of *q*ORAC: a surprisingly versatile and robust framework

Illustrations of the *q*ORAC framework are given in Figures 3, 4, S1–S3 (full details and code of simulations may be found in the SI).

A simple network, with two inputs and one output, is shown in Figure 3. The network requires two internal sensor metabolites, chosen to be the ones nearest the external input concentrations. Upon changes in external concentrations, the sensor concentrations change, causing changes in enzyme synthesis, which finally result in adaptation to the new optimum. The enzyme synthesis relations are also illustrated. Note the simplicity of these functional relationships, suggesting that simple gene networks could be constructed that can approximate them well. To illustrate the general applicability of *q*ORAC, consider the complicated branched example network in Figure S1. It has two inputs and two outputs and two allosteric interactions; by employing four sensors, it can be made robust to changes in all four external concentrations.

**Figure 3.**
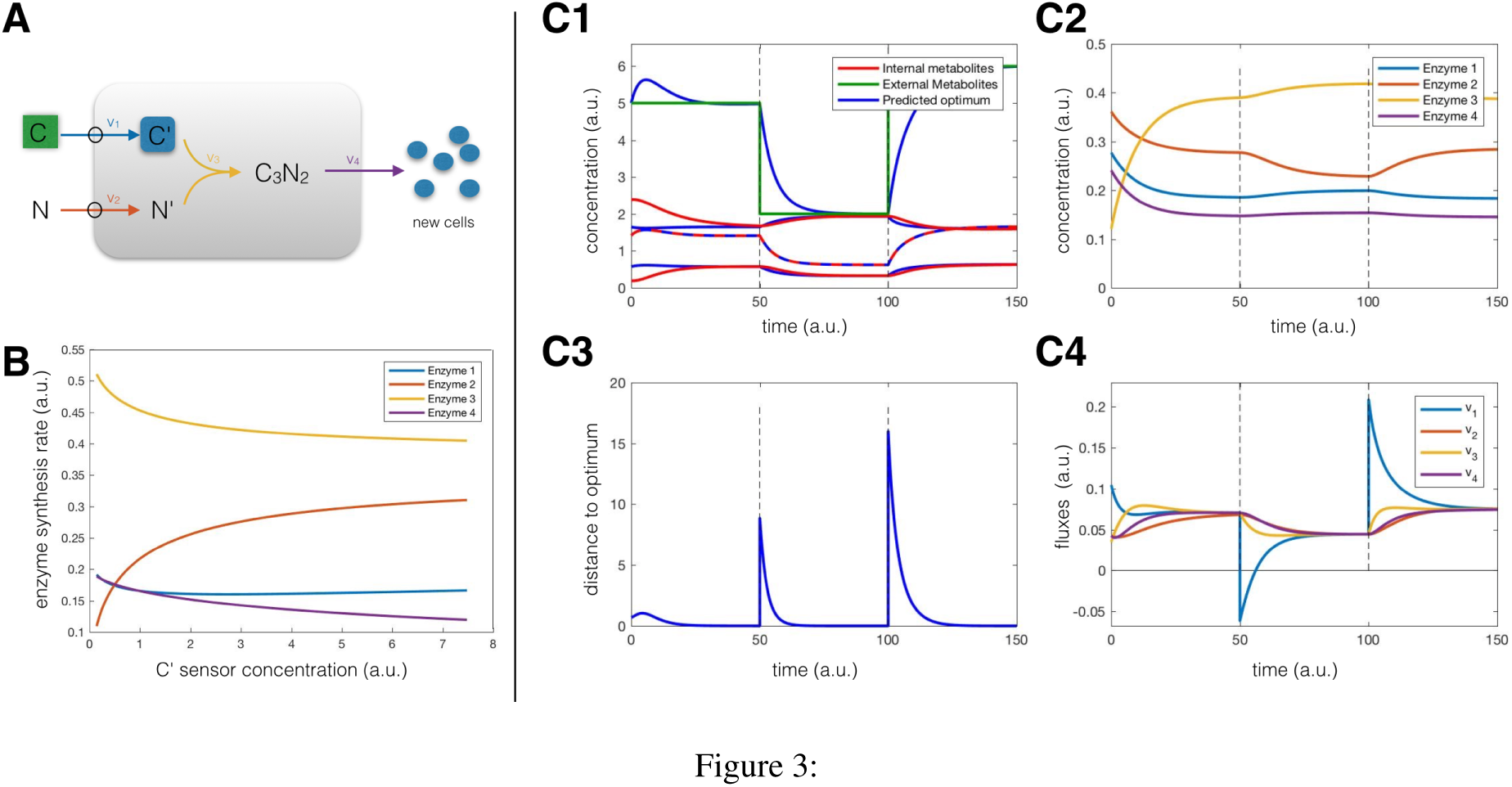
Example *q*ORAC dynamics. The dynamics are illustrated for the network shown in A. The green box depicts a varying external concentration, the blue box denotes the sensor concentration. B: the optimal input-output relations, showing enzyme synthesis rates as a function of changing sensor concentration *C*′. Inplots C1 to C4, the external *C* concentration is changed after 50 time units, and again after 100 time units. C1: The optimum predicted by the sensor (blue lines) converges to dynamic metabolite concentrations (red) and fixed external metabolite concentration (green), even when the external concentration changes at *t* = 50 and *t =* 100. The sensor is shown in dashed red-blue. C2: enzyme dynamics equilibrate after each change in external conditions, and reach their optimal levels. C3: the steered metabolic pathway reaches the optimum after each external change, as the distance to the (periodically changing) optimum reaches zero after some time. C4: flux dynamics equilibrate, showing that the pathway has reached steady state each time the external conditions change. Full equations, parameters and code are given in the SI.

The *q*ORAC control does not guarantee that a metabolic pathway is actually steered towards the optimum. In an example in which one of the periodically changing parameters is a *K_m_* parameter of a rate law, the choice of sensors matters critically (Figure 4). With one choice, the system robustly steers to the optimal specific flux steady state, but with another choice it does not. In both cases, the technical requirements to use the internal metabolites as sensors are met.

**Figure 4.**
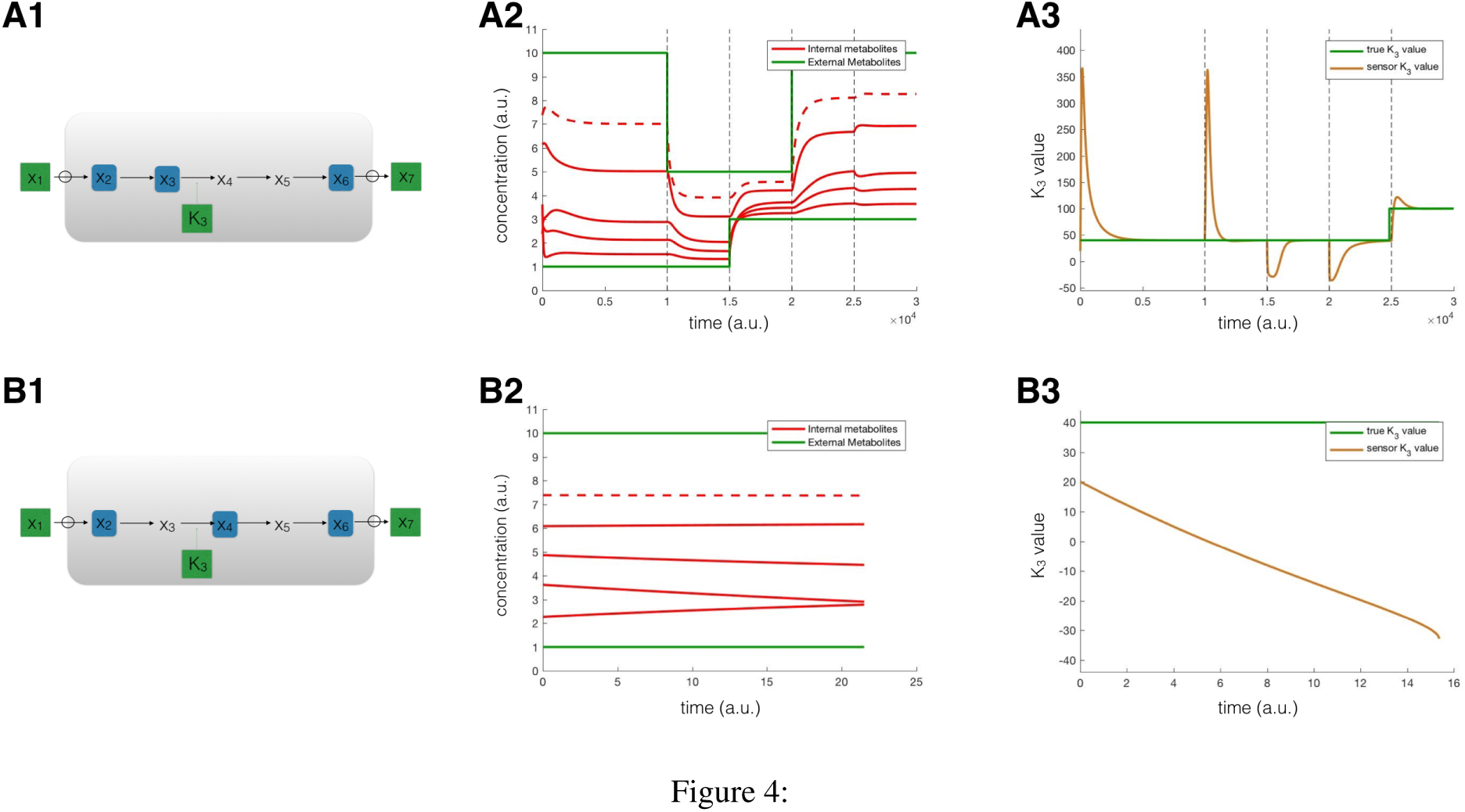
*q*ORAC for an internal parameter. In this example *q*ORAC is illustrated for a *K_m_* parameter in the third reaction, *K*_3_. In A1 and B1, the same pathway is drawn, but with different choices of sensors (in blue). A2: metabolite dynamics in which first external concentrations are varied (green) and at the end also *K_3_* is varied. A3: *K_3_* (in green) is varied at time *t* = 2500, and the predicted optimal value (in orange) subsequently converges, illustrating robust adaptive control. In B1, sensor *x_3_* is swapped with *x*_4_. The dynamics of metabolites (B2) and predicted *K_3_* values (B3) do start to change. However, the dynamics converge to a singular point, and the dynamical system can not continue. This second choice of sensors does not yield a gene expression control system which steers the pathway to optimal specific flux.

The *q*ORAC framework is able to start from nearly any initial condition. As an extreme example, with no enzymes present, and only the sensor concentration and no other internal metabolite, the *q*ORAC-controlled pathway still steers to optimum (Figure S2). Similarly, if the sensor concentrations are ‘wrong’, such that they predict a metabolic flow in the opposite direction to the one dictated by external concentrations, the combined controlled system nevertheless converges to the correct optimum (Figure S3).

### Concrete biological examples

In each of the pathways introduced in Figure 2A-E, the sensor metabolite(s) and transcription factor(s) have been identified. Specifying the kinetics for each enzymatic step in the pathway now directly gives the corresponding objective function (**7**) and the *q*ORAC framework can be set up. In two of these examples there is previous work on optimal metabolic rates, so we consider these in more detail.

The case of galactose uptake (Figure 2B) in yeast has been studied theoretically in detail by Berkhout et al.(2013), including fitting the parameters in the experimentally known gene network to approximate the theoretically predicted optimal input-output relations. Recent experimental evidence moreover shows that yeast cells are indeed able to tune the levels of these enzymes to optimise growth rate (Figure 1A).

For optimal ribosomal synthesis (Figure 2D), there is also previous theoretical work related to optimality. Maximal growth rates in *E. coli* are closely related to the expression of optimal concentrations of ribosomes: the translation machinery, including ribosomes, forms the largest protein fraction at maximal growth rate (Li et al., 2014), and is therefore most likely under tight control. Two recent models were able to reproduce the optimal synthesis of ribosomal concentration to maximise biomass synthesis rate (Scott et al., 2014; Bosdriesz et al., 2015). The model by Scott et al. (2014)may be viewed as a direct phenomenological implementation of *q*ORAC for a linear chain

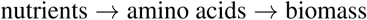

in which the amino acid concentrations are used as sensor. In the Bosdriesz et al. (2015)model, there is more biological detail. The main difference between our approach and (Scott et al., 2014;Bosdriesz et al., 2015) is that in *q*ORAC the input-output relations are predicted directly and only from the structure and kinetics of the pathway, rather than being chosen by the modellers. The qualitative nature of the input-output relations is of course identical.

## Discussion

Experimental evidence is accumulating that metabolic pathways are indeed able to optimise their enzyme resources to maximise growth rate. We addressed what type of enzyme synthesis rates, influenced by metabolite-binding transcriptions factors, result in maximisation of specific fluxes in metabolic pathways, a requirement for growth rate maximisation. We demanded robustness of optimality in the face of environmental changes. We presented the *q*ORAC framework, which implements such control for Elementary Flux Modes, the minimal steady state pathways that maximise specific flux (Müller et al., 2014;Wortel et al., 2014).

We use the term Specific Flux (g) Optimisation by Robust Adaptive Control (*q*ORAC) to describe the regulatory mechanism. ‘Robust’ signifies that attaining optimal states is independent of (environmental) parameter values – the system is robust to them. ‘Adaptive’ means that the control system steers the metabolic system to optimality without direct knowledge of external changes, contrary to the more widely studied problem of ‘optimal control’, in which the steering mechanism works using external changes as inputs to the controller (Krstic et al., 1995).

An important finding of our work is that the number of sensor metabolites must be (at least) equal to the number of parameters for which the metabolic pathway is robustly optimal. In other words, if the metabolic pathway always achieves states of maximal specific flux regardless of the values of three (independently-changing) environmental parameters, such as, for example, osmolarity, temperature and some nutrient concentration, then the number of sensors is expected to be three. This is a general result that follows from the associated mathematics of this control problem. Finding the sensors experimentally is difficult, and the number of known sensors is still quite small. However, it is telling that the whole of central metabolism in *E. coli* seems to be controlled by just three sensors, FBP, cAMP and F1P (Kochanowski et al., 2017).

The location of suitable sensors does not follow immediately from the optimisation problem. In general, one needs to make sure that the Implicit Function Theorem applies to the optimum equations (**9**), and this is not a trivial matter. However, a different argument shows that sensors near the beginning or ends of the pathway would work in most cases. The reason is that for all metabolites in between a set of fixed concentrations, their optimal value is uniquely determined by minimising the corresponding optimisation problem (i.e. finding the minimum of a suitable objective function *O*(*x; x*_S_) with *x* the set of metabolites between the sensors *x*_S_). The remaining variables, including the external concentrations, then need to be determined using the optimum equations (**17**). This is easiest exactly when the sensor is close to the external metabolite. Also from a biological standpoint this makes sense: such sensors obviously provide the most information of any change in external concentrations.

An important question is whether the adaptive control can be achieved by molecular circuits, given our understanding of biochemical kinetics and molecular interactions. The explicit example from galactose metabolism in yeast (Berkhout et al., 2013) gives hope that this might be true in general. If the necessary gene network is small, then the optimal circuit is likely also evolvable. We cannot give definite answers about this, but the computational analyses of different circuits, of which some are shown in this paper, indicate that optimal circuits show remarkably simple dynamics and input-output relations. One would expect that biochemical systems are capable of evolving those, and that synthetic biologists are capable of designing them.

The parameterisation of the optimising circuit is completely determined by the kinetics and the wiring of the metabolic pathway that it controls, since the objective function (**7**) contains all this information. This interdependence between the controller and the controlled is sometimes called the ‘internal model principle’ in engineering (Francis and Wonham, 1976) which roughly states that the control system should have knowledge of the dynamic behaviour of the system in order to be able to control it. Additional control mechanisms may then prevent for instance undesired oscillations or slow responses.

Technological advances have spurred recent interest in studying control properties of gene regulatory networks in cellular metabolism. One line of work involves characterising a particular gene control system and studying its theoretical properties. Examples are the perfect adaptation in the chemotaxis network in *E. coli* (Barkai and Leibler, 1997;Yi et al., 2000), the robustness properties of the heat-shock response system (El-Shamad et al., 2005) and of the circadian clock (Stelling et al., 2004). Several authors have considered dynamic optimisation of resources in pathways from a mostly computational perspective, e.g. to minimise the time of adaptive response (Pavlov and Ehrenberg, 2013), deFBA (Waldherr et al., 2015), and for other objectives than maximal specific flux, such as detecting equilibrium regimes of pathways (Oyarzún et al., 2012), robustness to flux perturbations (Oyarzún and Stan, 2012), and noise propagation (Oyarzún et al., 2015). In many studies, the control is not adaptive, but optimal; the objective is then usually to maximise the long term production of biomass (van den Berg et al., 1998;Pavlov and Ehrenberg, 2013;Giordano et al., 2016, e.g.).

The approach taken here differs principally from previous works in the following respect. The objective (maximal specific flux) is defined in advance, and the optimal input-output relations are characterised later. The framework is also analytic rather than computational: the input-output relations are obtained by solving the optimum equations (**9**) for the pathway under study, rather than by using an optimisation routine.

The choice of sensors sometimes matters for the control to steer the pathway to optimum (Figure 4). This example already indicates that, although the *q*ORAC control follows logically from the design objective, it is not easy to decide which intermediate metabolites make it controllable. We cannot expect completely general mathematical theorems. Apparently, some choices of sensors do work, and others do not, for the same pathway, using the same initial conditions.

*q*ORAC has direct applications in synthetic biology. To achieve maximal production rates in a biotechnological-product producing pathway requires a controller that *q*ORAC provides. The only ingredient to design such a controller are the enzymatic rate laws in the pathway. *q*ORAC then immediately makes predictions about the optimal enzyme synthesis rates, as a function of one or more intermediate metabolites. As the synthetic biology field advances, synthetic circuits with the required input-output relationships for the constituent enzymes of the pathway can be designed and built. *q*ORAC therefore does not only contribute to the general understanding of steering mechanisms to optimal states, but provides direct operational relevance for microbiology, synthetic biology and biotechnological applications.

## Author contributions

R.P., B.T. and F.J.B. designed the research; R.P., J. Hulshof, J. Hendriks, B.T. and F.J.B. performed the research; R.P. wrote the numerical code and performed the simulations; R.P., B.T. and F.J.B. wrote the paper; R.P. and J. Hendriks wrote the SI.

## Acknowledgements

F.J.B. acknowledges funding of Nederlandse Organisatie NWO-VIDI project No. 86411-011; B.T. acknowledges funding of Nederlandse Organisatie NWO-VICI project 865.14.005.

The authors declare no conflict of interest.

